# VCF2CNA: A tool for efficiently detecting copy-number alterations in VCF genotype data

**DOI:** 10.1101/131235

**Authors:** Daniel K. Putnam, Ma Xiaotu, Stephen V. Rice, Yu Liu, Jinghui Zhang, Xiang Chen

## Abstract

VCF2CNA is a web interface tool for copy-number alteration (CNA) analysis of VCF and other variant file formats. We applied it to 46 adult glioblastoma and 146 pediatric neuroblastoma samples sequenced by Illumina and Complete Genomics (CGI) platforms respectively. VCF2CNA was highly consistent with a state-of-the-art algorithm using raw sequencing data (mean F1-score=0.994) in high-quality glioblastoma samples and was robust to uneven coverage introduced by library artifacts. In the neuroblastoma set, VCF2CNA identified MYCN high-level amplifications in 31 of 32 clinically validated samples compared to 15 found by CGI’s HMM-based CNA model. The findings suggest that VCF2CNA is an accurate, efficient and platform-independent tool for CNA analyses without accessing raw sequence data.

## Background

Copy-number alterations (CNAs) are gains or losses in chromosomal segments that frequently occur in tumor cells. Recent surveys suggest that certain cancers are driven by CNAs [1]. In addition to directly affecting cancer genes (e.g., *MYCN* and *MDM2* amplifications and *RB1* and *CDKN2A* deletions), CNAs mediate oncogene overexpression through enhancer hijacking [2-5]. Several experimental methods are available to identify CNAs in tumor cells. Fluorescence in situ hybridization provides direct evidence of CNAs and is the gold standard for CNA detection in a targeted region [6]. Before the development of next-generation sequencing (NGS) technologies, array comparative genomic hybridization and high-resolution single nucleotide polymorphism (SNP) arrays permitted genome-wide evaluation of CNAs at 30-kb to 100-kb resolution.

The development of NGS, especially whole-genome sequencing (WGS) platforms, has revolutionized the detection of somatic mutations, including CNAs, in cancer samples. For example, Copy Number Segmentation by Regression Tree in Next Generation Sequencing (CONSERTING) [7] incorporates read-depth and structural-variation data from BAM files for accurate CNA detection in high-coverage WGS data. However, CONSERTING and other WGS-based CNA algorithms produce a fractured genome pattern (i.e., a hypersegmented CNA profile with an excessive number of intrachromosomal translocations) in samples with library construction artifacts [7], which poses a major challenge for precise CNA inference. Our extensive analysis indicated that although CNA and structural-variation detection was severely impaired by library artifacts, point-mutation detection was largely unaffected (data not shown), suggesting that a robust CNA tool can be developed from the variant information. Moreover, CONSERTING and other WGS algorithms require direct access to aligned BAM files. Most algorithms further incur complicated installation steps, which create barriers for their widespread adoption. Advances in technology and declines in costs have made NGS a commodity for both basic research and clinical service. Therefore, a robust CNA analytical tool that is efficient, convenient, and robust to library artifacts is needed to manage the demands of NGS data analysis.

VCF2CNA (http://vcf2cna.stjude.org) is a web-based tool for CNA analysis. The preferred input to VCF2CNA is a Variant Call Format (VCF) file. VCF is a widely adopted format for genetic variation data exchange, and VCF files are quite small compared to WGS BAM files. Each variant in a typical VCF file contains its chromosome position, reference/alternative alleles, and corresponding allele counts, which are used by VCF2CNA to identify copy-number alterations. This tool also accepts input in the Mutation Annotation Format (MAF) and the variant file format produced by the Bambino program [8].

## Results

VCF2CNA has a simple interface (Fig. 1a). The sole input is a VCF file (or a file in one of the other supported variant file formats) from a paired tumor–normal WGS analysis, which is uploaded via the interface to a web server where the application runs. The results are returned to a user-provided email address. VCF2CNA consists of two main modules: 1) SNP information retrieval and processing from the input data and 2) recursive partitioning–based segmentation using SNP allele counts (Fig. 1b). Actual running time for a typical sample is approximately 30 to 60 minutes, depending on the complexity of the genome.

**Fig. 1.**
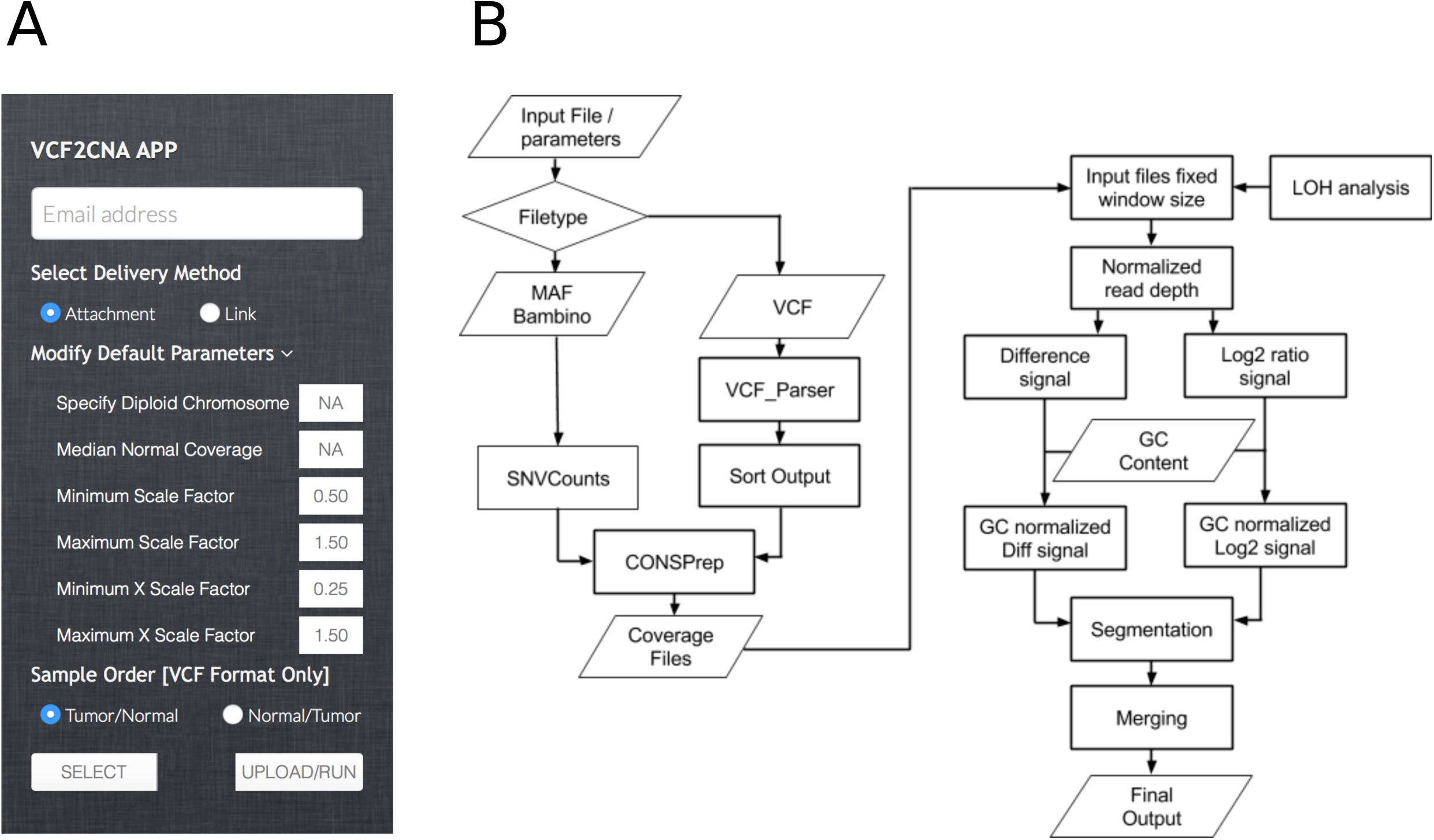
Overview of the VCF2CNA process. **a** User interface with parameters. **b** Server side pipeline. A parallelogram depicts input or output files, a rectangle depicts an analytical process, and a diamond depicts the condition for a follow-up process.

To evaluate the utility of VCF2CNA, we ran it on 192 tumor–normal WGS data sets. These sequences comprised 46 adult glioblastomas (GBMs) from The Cancer Genome Atlas (TCGA-GBM) dataset [9], sequenced by Illumina technology, and 146 pediatric neuroblastomas (NBLs) from the Therapeutically Applicable Research to Generate Effective Treatments (TARGET-NBL) dataset (unpublished), sequenced by Complete Genomics, Inc. (CGI) technology. On average, VCF2CNA used approximately 2.8 million high-quality SNPs per sample (median 2,811,245; range, 2,029,467–3,519,454 in TARGET-NBL data) to derive CNA profiles.

### CNA analysis of TCGA-GBM data

The adult TCGA-GBM data downloaded from dbGaP (accession number: phs000178.v8.p7) included 46 samples. We first evaluated VCF2CNA’s resistance to library construction artifacts by using 24 samples from this set, which were previously identified as having a fractured genome pattern by CONSERTING and other CNA algorithms [7]. Indeed, VCF2CNA produced CNA profiles that are globally consistent with those of SNP array–derived CNA profiles (downloaded from TCGA, Additional file 1.1 and 1.2) and more robust to noise than those produced by CONSERTING. Specifically, VCF2CNA yielded a mean 59.4-fold reduction in the number of predicted segments than did CONSERTING (median, 46.2; range, 16.2–285.7; *p*□= □3.0 × 10^−6^ by Wilcoxon signed-rank test, Fig. 2a and Additional file 1).

**Fig. 2.**
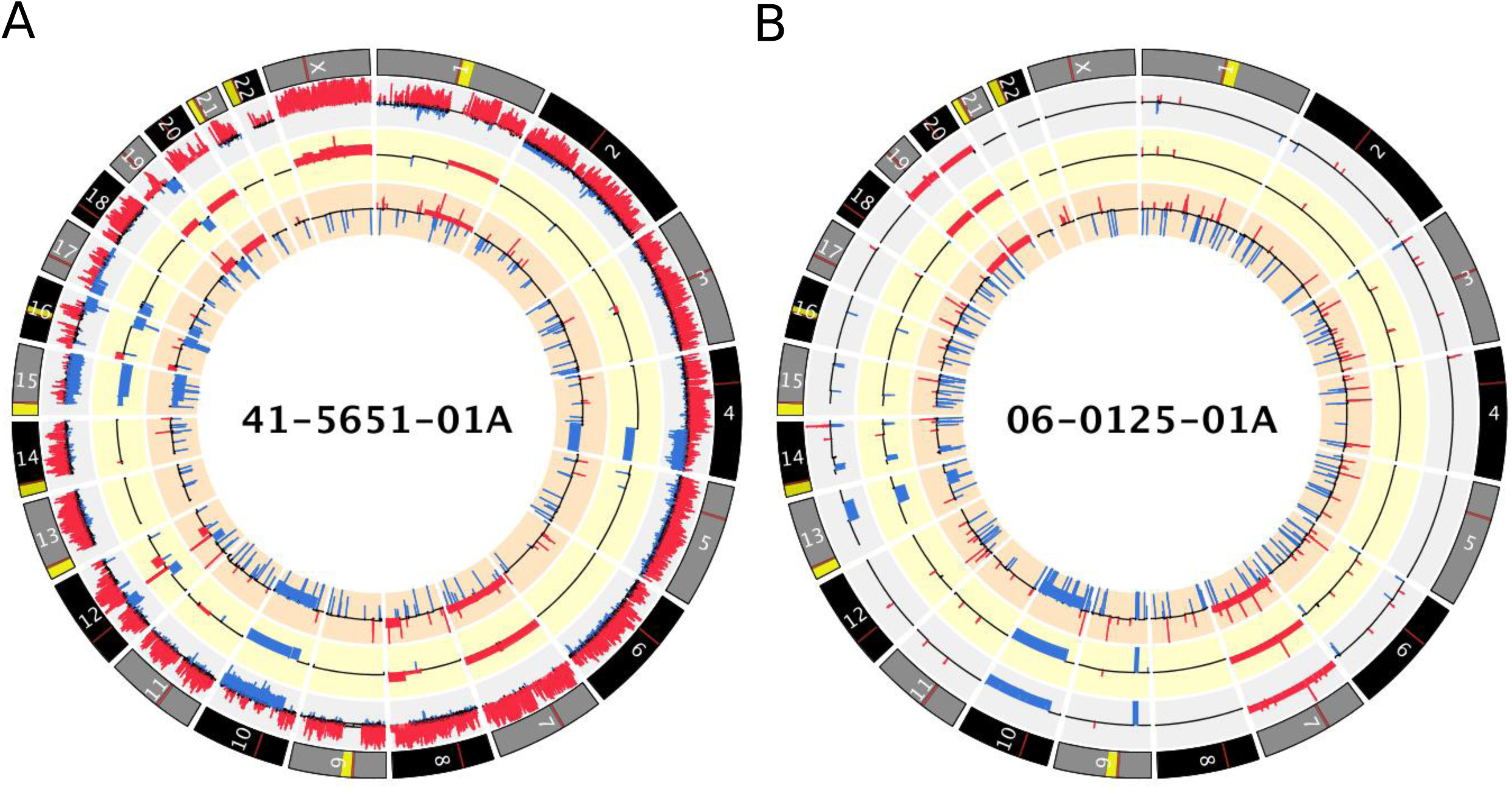
A Circos plot that displays CNAs found by CONSERTING (outer ring), VCF2CNA (middle ring), and SNP array (inner ring) for **a** TCGA-GBM fractured sample 41-5651-01A and **b** TCGA-GBM unfractured sample 06-0125-01A. Alternating gray and black chromosomes are used for contrast. Yellow regions depict sequencing gaps, whereas red regions depict centromere location. Blue segments depict copy-number loss, and red segments indicate copy-number gain.

We used an F_1_ scoring metric [10] to measure the consistency between the CNA profiles derived from VCF2CNA and CONSERTING in the remaining 22 high-quality sample pairs (Fig. 2b and Additional file 2). These programs identified approximately 700 Mb of the CNA regions in each sample (range, 92–2299 Mb) with high consistency (mean F_1_ score, 0.9941; range, 0.9699–0.9995) (Table 1).

**Table 1.** F_1_ score between CONSERTING and VCF2CNA and autosomal CNAs per sample in 22 TCGA samples

| Sample | $F_1$ score | Autosomal CNAs per sample (Mb) |
| --- | --- | --- |
| SJHGG011906_D1_G1_N13 | 0.9699 | 567.70 |
| SJHGG010485_D1_G1 | 0.9840 | 789.90 |
| SJHGG011903_D1_G1 | 0.9862 | 459.20 |
| SJHGG010643_D1_G1_N5 | 0.9870 | 1471.67 |
| SJHGG010641_D1_G1 | 0.9884 | 285.89 |
| SJHGG010600_R1_G1 | 0.9892 | 485.85 |
| SJHGG010484_R1_G1_N2 | 0.9949 | 2299.48 |
| SJHGG010560_R1_G1 | 0.9955 | 756.08 |
| SJHGG010624_R1_G1 | 0.9956 | 1259.68 |
| SJHGG010600_D1_G1 | 0.9968 | 389.60 |
| SJHGG010485_R1_G1 | 0.9970 | 92.16 |
| SJHGG011904_D1_G1 | 0.9979 | 696.48 |
| SJHGG010540_D2_G1 | 0.9981 | 660.74 |
| SJHGG010484_D1_G1 | 0.9983 | 841.72 |
| SJHGG010509_D1_G1 | 0.9983 | 586.18 |
| SJHGG010560_D1_G1 | 0.9984 | 551.73 |
| SJHGG010577_D1_G1 | 0.9984 | 831.67 |
| SJHGG010509_R1_G1 | 0.9988 | 562.44 |
| SJHGG010572_R1_G1 | 0.9992 | 427.91 |
| SJHGG010572_D1_G1 | 0.9994 | 456.27 |
| SJHGG010624_D1_G1 | 0.9995 | 454.09 |
| SJHGG010540_R1_G1 | 0.9995 | 463.89 |

We evaluated the segmental overlap between the CONSERTING outputs and the VCF2CNA outputs for each sample. A CNA segment detected by CONSERTING was classified as corroborated if 90% of the bases in the segment received the same type of CNA call from VCF2CNA (Table 2). The comparison shows that VCF2CNA faithfully recapitulated medium to large CNA segments (≥ □100 kb) (Fig. 3a), whereas CONSERTING had greater power for identifying focal (< □100 kb) low-amplitude (absolute log2 ratio change < □1.0) CNAs (*p*□= □1.306 × 10^−5^ by Wilcoxon signed-rank test, Fig. 3b). Furthermore, the segmental–based analysis revealed that the detection power was less affected in focal CNAs with large amplitudes (log2 ratio□≥ □3.0) (Fig. 3c).

**Table 2.**
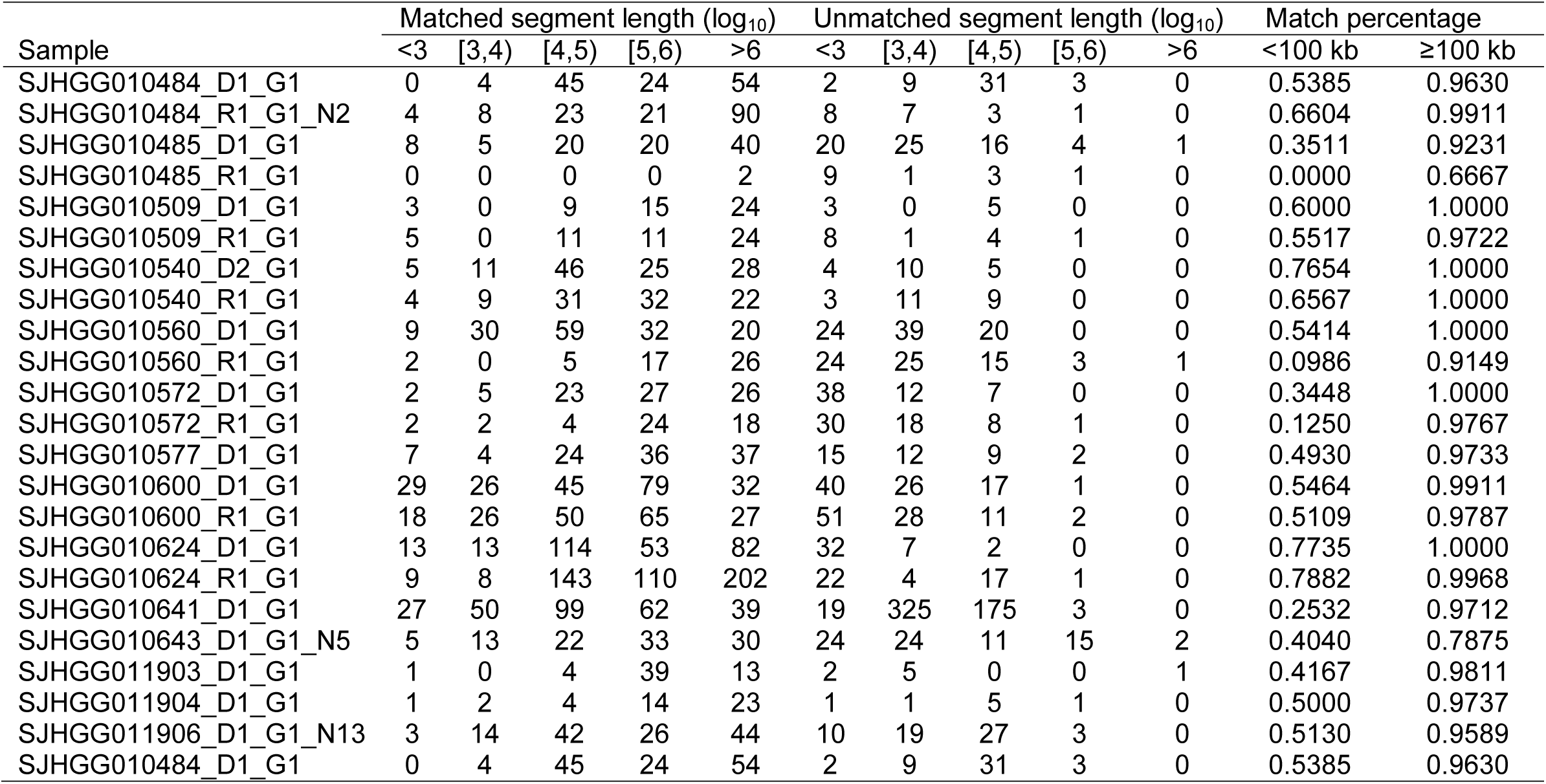
Counts of corroborated and uncorroborated segments by segment length

**Fig. 3.**
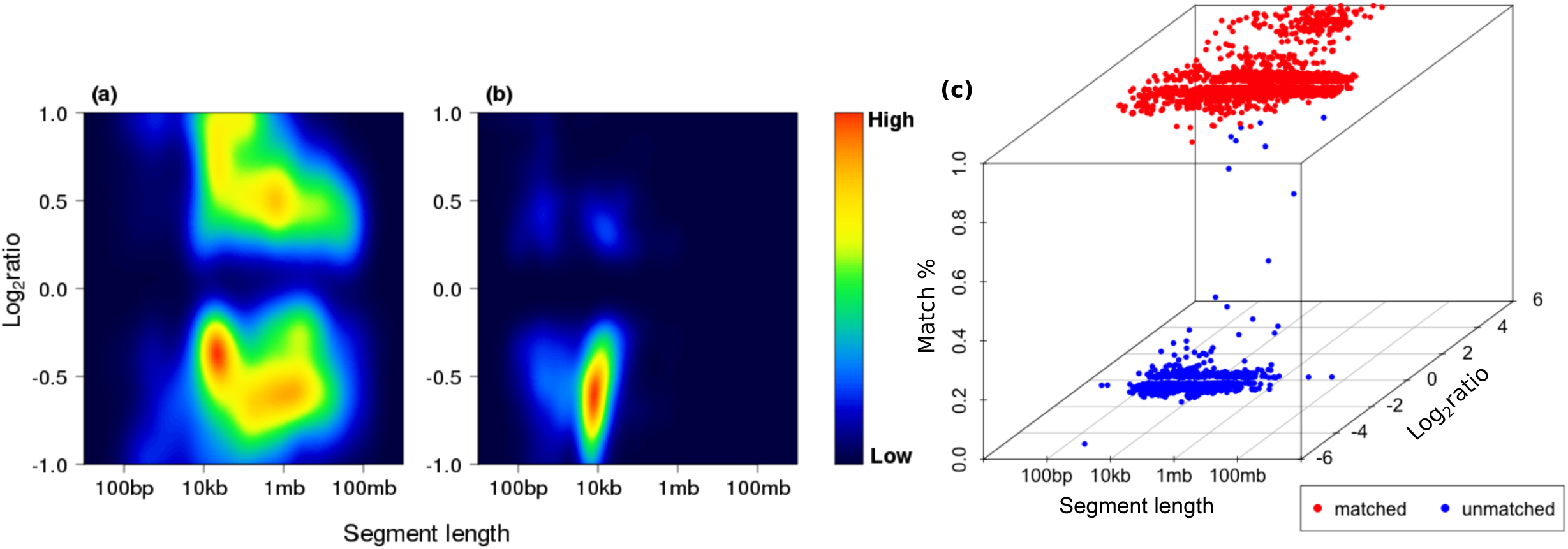
Heatmap of segment length by CNA intensity. Color scale depicts density of segment found at a given segment and CNA size. **a** Corroborated samples, **b** uncorroborated samples, and **c** three-dimensional plots of segment length, CNA intensity, and percent agreement with CONSERTING segments are shown.

To further test whether VCF2CNA accurately captures the CNA patterns in samples with library artifacts, we applied the cghMCR algorithm [11]. This algorithm identifies genomic regions that exhibit common gains and losses across all 46 samples from either VCF2CNA profiles or SNP array–derived CNA profiles (downloaded from TCGA). Although the signal from VCF2CNA contained less noise than did the signal from the SNP array in most samples (Additional file 1), both profiles reveal common recurrently amplified and/or lost regions (Fig. 4). These changes included chromosome-level changes (i.e., chr7 amplifications and loss of chr10) and segmental CNAs (i.e., focal deletion of the *CDKN2A/B* locus on chr9p) [12]. Moreover, VCF2CNA identified recurrent losses in *ERBB4* on chr2q and *GRIK2* on chr6q that were absent in the SNP array profiles. *ERBB4* encodes a transmembrane receptor kinase that is essential for neuronal development [13]. It is frequently mutated in patients with non-small cell lung cancer [14], and silencing of *ERBB4* through DNA hypermethylation is associated with poor prognosis in primary breast tumors [15]. Similarly, *GRIK2* is a candidate tumor suppressor gene that is frequently deleted in acute lymphocytic leukemia [16] and silenced by DNA hypermethylation in gastric cancer [17].

**Fig. 4.**
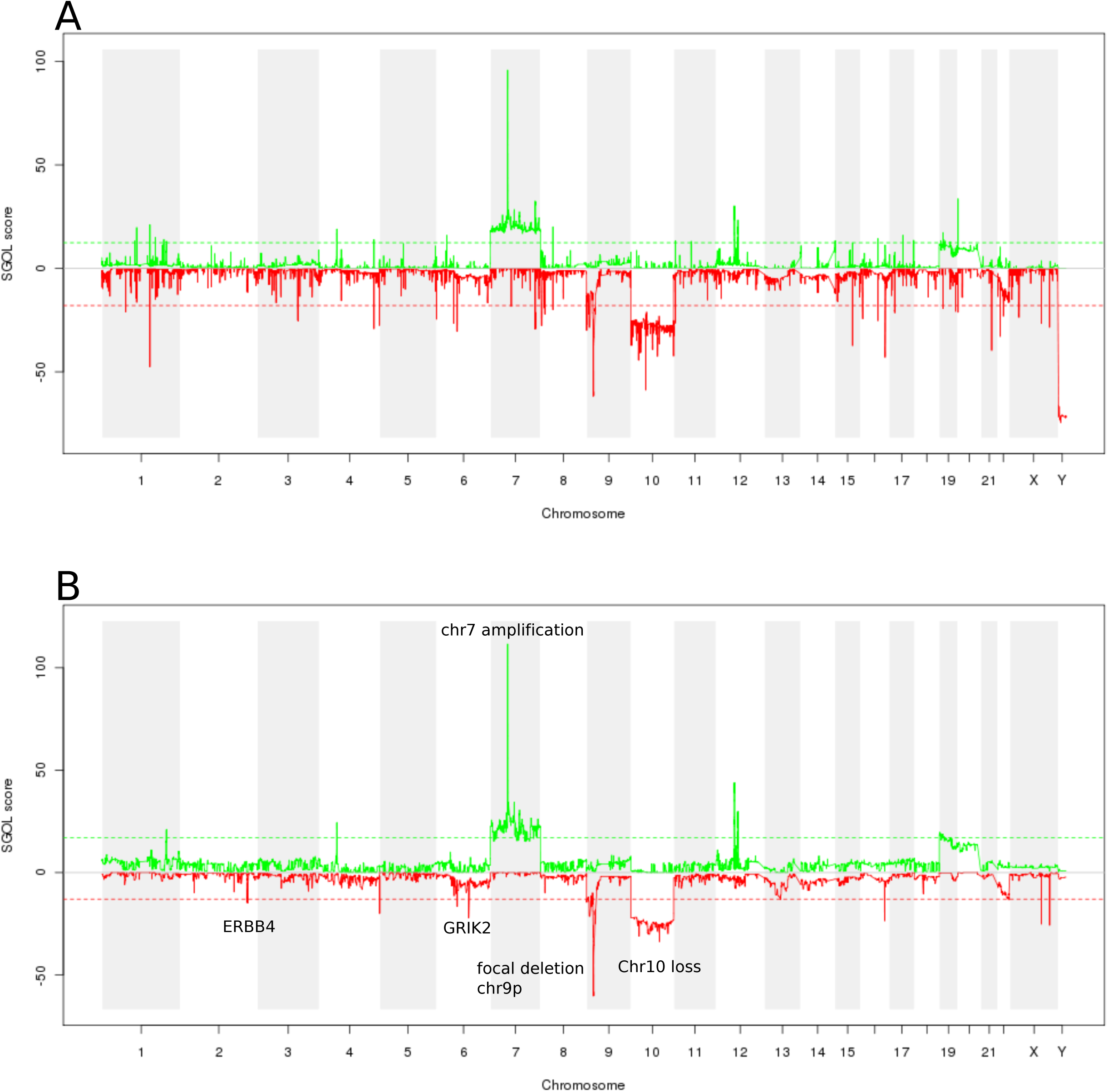
A chgMCR plot of 46 TCGA-GBM samples. **a** SNP array data and **b** VCF2CNA data are shown.

Amplifications such as double minute chromosomes and homogeneously staining regions represent a common mechanism of oncogene overexpression in tumors [18]. Among the 46 TCGA-GBM samples analyzed, VCF2CNA identified double minute chromosomes in 34 samples affecting the *EGFR* [19], *MDM2* [20], *MDM4* [21]*, PDGFRA* [22]*, HGF* [23]*, GLI1* [24]*, CDK4* [25], and *CDK6* [26] genes (Fig. 5 and Additional file 3). These events consisted of high-level amplifications in 21 samples with potential fractured genome patterns (Additional file 3a) and 13 previously reported samples (Additional file 3b) [7, 27].

**Fig. 5.**
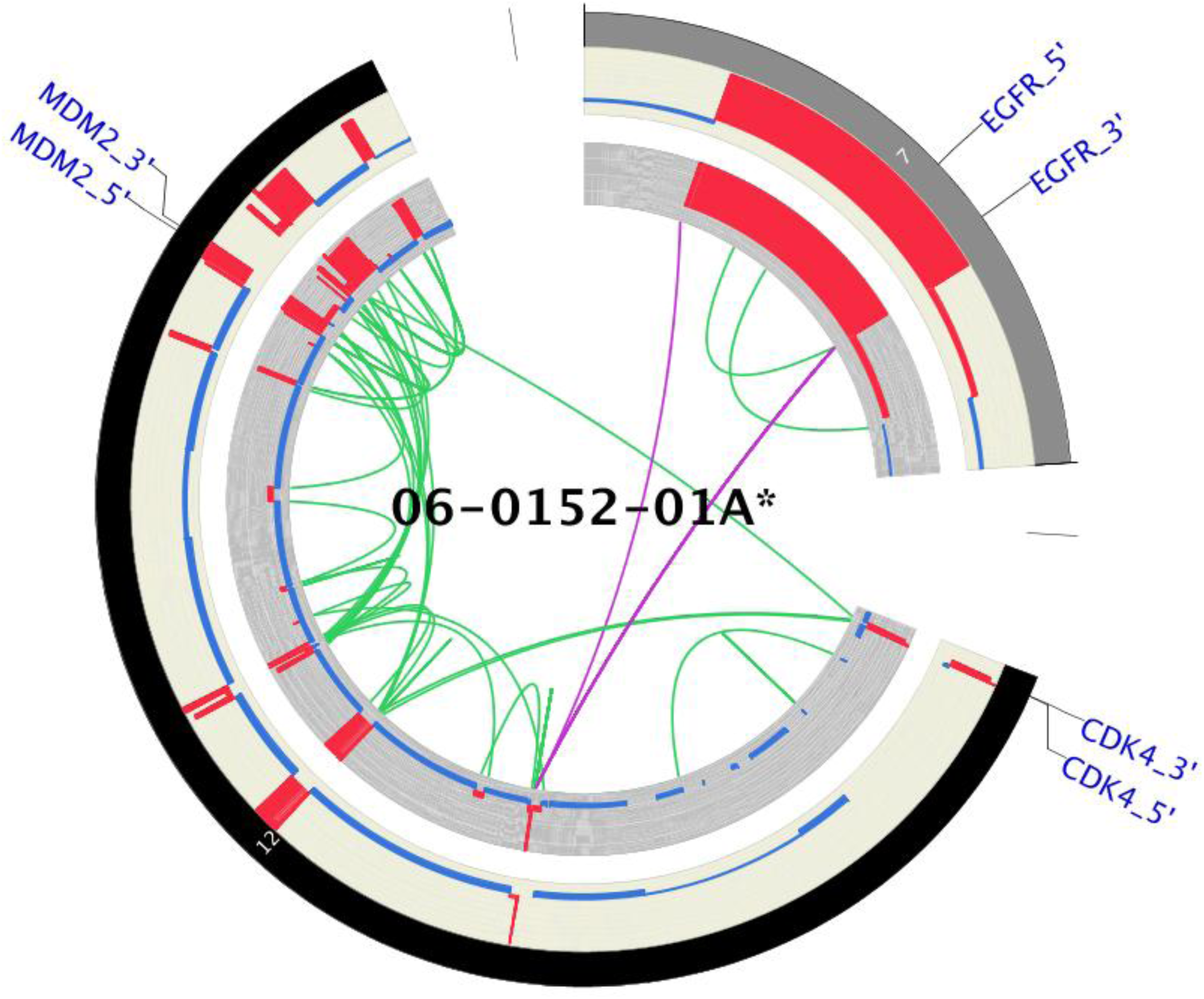
A Circos plot of VCF2CNA (outer ring) and CONSERTING (inner ring), depicting high-amplitude focal CNA segments in TCGA-GBM sample 06-0152-01A. Included in these segments are the known cancer genes *EGFR*, *CDK4*, and *MDM2*.

### CNA analysis of TARGET-NBL data

We applied VCF2CNA to the TARGET-NBL dataset downloaded from dbGap (assession number: phs000467). This dataset consists of 146 tumors with matched normal WGS samples, sequenced with CGI technology. Because the ligation-based CGI technology has notable differences in the detection of single nucleotide variants (SNVs) and insertions/deletions (indels) compared to Illumina systems [28], this dataset provided an opportunity to evaluate VCF2CNA’s robustness using different sequencing platforms.

We used VCF2CNA to perform cghMCR analysis with CNA profiles and observed a genome pattern similar to that reported for SNP array platforms (Fig. 6a) [29]. In addition to loss of large regions on chr1p, 3p, and 11q and a broad gain of chr17q, VCF2CNA found frequent focal amplifications of *MYCN* in NBL tumors and several potential cancer-related CNAs, including high-level amplifications of *CDK4* (1 tumor), and *ALK* (2 tumors) (Fig. 6b).

**Fig. 6.**
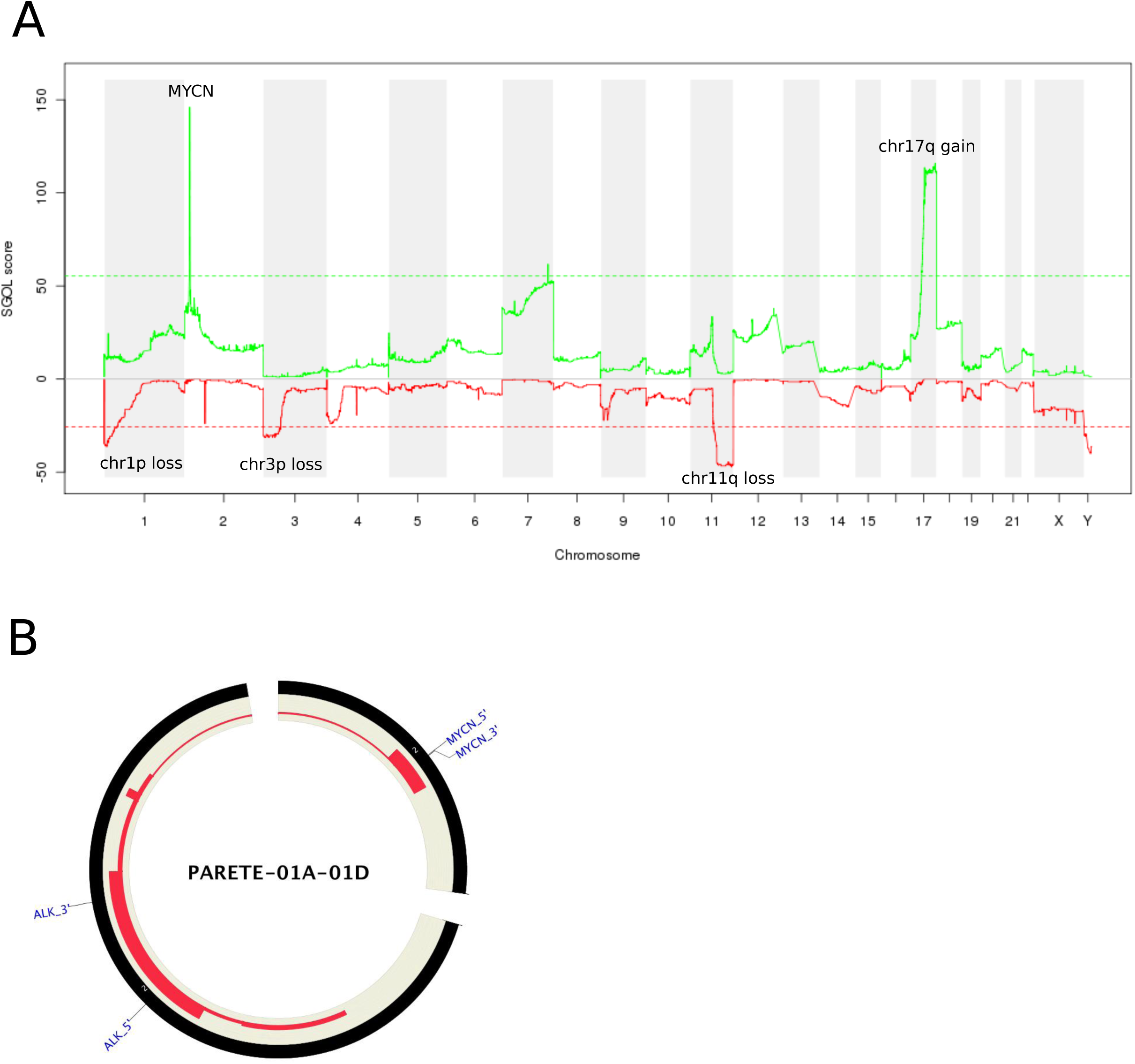
Analysis of the TARGET-NBL dataset, consisting of 146 tumors. **a** A chgMCR plot in which green depicts regions of copy-number gain and red depicts regions of copy-number loss. **b** A Circos plot showing a focal gain on chromosome 2 for *MYCN* and *ALK5* for sample PARETE-01A-01D.

High-level amplification of *MYCN* is a known oncogenic driver found in ∼25% of pediatric patients with NBL, and is associated with aggressive tumors and poor prognosis [30]. A subset of 32 tumors in the TARGET-NBL cohort contains clinically validated amplifications of *MYCN*. Although the CGI’s hidden Markov CNA model (unpublished) reported *MYCN* amplifications in 15 of these 32 tumors, VCF2CNA successfully identified high-level amplifications in 31 tumors. In the clinically validated *MYCN*-amplified sample that went undetected by VCF2CNA, a follow-up review revealed that tumor heterogeneity and sampling bias most likely contributed to the discrepancy. Moreover, VCF2CNA predicted two additional *MYCN* amplification events among the remaining tumor samples, indicating that VCF2CNA can identify clinically relevant CNAs that were undetected by traditional methods of CNA detection. The high-level concordance with clinically validated data provides a strong indication that VCF2CNA is applicable to multiple tumor types collected from different sequencing platforms.

### Discussion and conclusions

We developed VCF2CNA for the systematic and robust detection of CNAs from VCF and other genotyping variant call formats. Analysis of 192 paired tumor–normal WGS samples sequenced on multiple platforms demonstrates that VCF2CNA is robust to library construction artifacts and captures medium to large CNA segments with high accuracy. Because VCF2CNA is robust to library artifacts and is highly accurate, it identified recurrent losses in potential tumor suppressors that were undetectable by alternative approaches.

VCF2CNA was designed with SNPs that were (on average) thousands of base pairs apart, which limits support for identifying focal copy-number changes. Therefore, state-of-the-art CNA algorithms have superior detection power for focal low-amplitude CNAs in high-quality, high-coverage WGS data.

In conclusion, VCF2CNA is a web-based tool that is capable of accurate and efficient detection of CNAs from variants called from high-coverage WGS data sequenced on various platforms.

## Methods

### Server availability

VCF2CNA is available at https://vcf2cna.stjude.org.

### Parameter definitions

The Specify Diploid Chromosome parameter normalizes results by the specified chromosome. The Median Normal Coverage parameter permits input of the median coverage value of SNPs from normal samples. The Minimum Scale Factor (autosomes) parameter is multiplied by the median to compute the minimum coverage value. The Maximum Scale Factor (autosomes) parameter is multiplied by the median to compute the maximum coverage value. The Minimum X Scale Factor is the minimum scale factor for chromosome X. The Maximum X Scale Factor is the maximum scale factor for chromosome X. The Sample Order (VCF format only) parameter defines the ordering of tumor and normal samples. VCF inputs must include tumor and normal data after the FORMAT field. Selecting the Tumor/Normal button assigns the tumor data to the first field after FORMAT and normal data to the second field. The Normal/Tumor radio button specifies the reverse order.

### Input data for VCF2CNA

The input for VCF2CNA analysis includes VCF, MAF, and the variant file format produced by the Bambino program. A fixed window size of 100 bp is used to obtain the mean coverage for each window. Windows with no variants are ignored. The mean read depth per window can be normalized to a set of reference diploid chromosomal regions by using the same criteria as CONSERTING or specified via the Specify Diploid Chromosome parameter.

### Run-time analysis

Single VCF files must be converted to a paired tumor/normal file before uploading. Alternatively, VCF2CNA accepts MAF and Bambino variant file formats. After uploading files to the server, the median running time was 23 minutes on an intel Xeon E5-2680 processor at 2.70 Ghz with 64 GB RAM. Server processing occurs in two principal steps: 1) preprocessing and SNP information extraction from input files and 2) running the recursive partitioning segmentation.

### F_1_ scoring metric and segmental corroboration

A genomic position was assigned a corroborated CNA call if its computed CNA type (gain or loss) by VCF2CNA matched the call computed by CONSERTING. A CAN segment in the CONSERTING profile was corroborated in the VCF2CNA profile if ≥ □90% of the segment positions were corroborated. The F_1_ score is given by 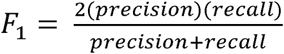. It was used to summarize the accuracy of VCF2CNA, compared with that of CONSERTING.

## VCF2CNA web server pipeline

### Step1 (snvcounts)

Single nucleotide variant frequencies are computed from the input file. For each chromosome and position, the values computed are TumorMutant, TumorTotal, NormalMutant, and NormalTotal. Additionally, the mean normal coverage is computed.

### Step2 (consprep)

The consprep program reads the SNV count data and incorporates a list of good / bad SNVs. It also reads a file specifying the number of 100-bp windows in each chromosome. If the total number of reads from the normal sample falls outside of the ranges specified by the options (median, minfactor, maxfactor, xminfactor, or xmaxfactor), the input position is ignored by the consprep step in the pipeline. The –xminfactor and – xmaxfactor settings apply to positions in chrX; the –minfactor and –maxfactor settings apply to all other chromosomes. The minimum coverage is the median multiplied by the –minfactor, and the maximum coverage is the median multiplied by the –maxfactor.

### Application

To run VCF2CNA, users should navigate to the application home page and click “run application.” The application runs on Google Chrome, Safari, Mozilla Firefox, and Microsoft Internet Explorer 11. Users must provide a valid email in the email address text field. Users will select whether results will be sent to the provided email address as either an email attachment or a link to the result files stored on the server. Results will be stored on the server for 14 days. Default run parameters may be modified depending on job specifications. Users should select the input file and click the “upload/run” button. The brower window should not be killed during the file upload. Once the file has been successfully uploaded, a notification will be displayed in the browser window and the user may discard the window.

### Rationale for not using the reciprocal-overlap rule

To compare CNA calls from different algorithms, the reciprocal 50% overlap criterion [28] is commonly used. This rule is not suitable when two CNA calls are derived from platforms with different powers in detecting focal CNAs. A considerably larger average distance occurred between adjacent probes. VCF2CNA-derived CNA calls have an inherently lower resolution than does CONSERTING. When a focal CNA identified through CONSERTING occurs on top of a large CNA fragment, CONSERTING breaks the region into multiple segments. Although the CNA fragments in the region are largely corroborated between the two CNA callers, potentially none of these fragments satisfied the rule of reciprocal 50% overlap (Additional file 4).

## Declarations

### Ethics approval and consent to participate

Not applicable.

### Consent for publication

Not applicable.

### Availability of data and material

Both datasets were downloaded from dbGaP (https://dbgap.ncbi.nlm.nih.gov). The TCGA-GBM data were downloaded from dbGaP (accession number: phs000178.v8.p7) and included 46 samples. The TARGET-NBL data were downloaded from dbGap (accession number: phs000467) and included 146 samples. VCF2CNA is available at https://vcf2cna.stjude.org.

### Authors’ contributions

JZ and XC conceived the concept. DP, XM, and XC designed the VCF2CNA algorithm. DP, XM, SR, and XC implemented the algorithm. DP, XM, YL, and XC performed the analysis. DP and XC wrote the manuscript.

## Acknowledgements

We thank Dr. Nisha Badders for editing the manuscript and Soubhadra Datta for server support.

## Additional files

**Additional file 1:** Circos plot of CONSERTING (outer ring), VCF2CNA (middle ring), and SNP array (inner ring) for 24 TCGA-GBM samples with a fractured gene signature.

**Additional file 2:** Circos plot of CONSERTING (outer ring) and VCF2CNA (inner ring) for all 22 TCGA-GBM samples without a fractured gene signature.

**Additional file 3:** A Circos plot of VCF2CNA (outer ring) and CONSERTING (inner ring), depicting high-amplitude focal CNA segments in 34 TCGA-GBM samples. **a** 21 fractured genome TCGA-GBM samples. **b** 13 previously reported samples.

**Additional file 4:** Segmental Overlap. **a** A hypothetical large segment identified by CONSERTING (red). **b** Subsequent focal segments identified by CONSERTING (blue). The original segment was split into five subsegments. None of the subsegments in b met the reciprocal 50% segment overlap criteria with the original segment.

